# pH-mediated inhibition of a bumble bee parasite by an intestinal symbiont

**DOI:** 10.1101/336347

**Authors:** Evan C Palmer-Young, Thomas R Raffel, Quinn S McFrederick

**Affiliations:** Department of Entomology, University of California Riverside, Riverside, CA, USA; Department of Biology, Oakland University, Rochester, MI, USA

## Abstract

Non-pathogenic microbes can provide multiple benefits to their hosts, including pathogen inhibition. Gut symbionts can augment resistance to pathogens by stimulating host immune responses, competing for space and nutrients, or producing antimicrobial metabolites. The gut microbiota of social bees, which pollinate many crops and wildflowers, has demonstrated benefits against diverse infections and might help protect against pathogen-related declines. The bumble bee gut microbiota, consisting chiefly of five taxa common to corbiculate bees, has been shown to enhance resistance to the trypanosomatid parasite *Crithidia bombi*. Specifically, infection intensity was negatively correlated with abundance of *Lactobacillus* “Firm-5” bacteria. However, the mechanism underlying this relationship remains unknown. We tested the hypothesis that the Firm-5 bacterium *Lactobacillus bombicola*, which produces lactic acid, inhibits *C. bombi* via a pH-mediated effect.

Consistent with our hypothesis, *Lactobacillus bombicola* spent medium inhibited *C. bombi* growth via a reduction in pH that was both necessary and sufficient for inhibition. Inhibition of all parasite strains occurred within the pH range previously observed in honey bee guts, though sensitivity to acidity varied among parasite strains. Spent medium was slightly more potent than HCl, D-, and L-lactic acids for a given pH, suggesting that other metabolites also contribute to inhibitory effects. Our results implicate symbiont-mediated reduction in gut pH as a key determinant of trypanosomatid infection in bees. Future investigation into *in vivo* effects of gut microbial composition on pH and infection intensity would help determine the relevance of these findings for bees threatened by trypanosomatids.

**Importance:** Pollinators such as honey and bumble bees provide services to plants in agricultural and wild ecosystems, but both wild and managed bees are threatened by infection-related declines. The symbiotic gut microbiota of bees provides a naturally occurring defense against infection. For example, the bumble bee microbiota reduces infection with trypanosomatid parasites, but how inhibition occurs remains unknown. We show that the acidic spent medium from a common bumble bee gut symbiont, *Lactobacillus bombicola,* inhibits *in vitro* growth of the trypanosomatid gut parasite, *Crithidia bombi.* The acidity of the spent medium was both necessary and sufficient for parasite inhibition. Inhibitory pH values were within the range documented in honey bee guts, suggesting that pH-mediated parasite inhibition is plausible in live bees. Results suggest that production of acids by sugar-fermenting symbionts confers pH-mediated resistance to infection in bees, whereas depletion of core microbiota could result in low-acid conditions that favor parasite growth.

## INTRODUCTION

Both animals and plants associate with symbiotic bacterial communities that provide functional benefits to their hosts (1). The gut microbiota comprise some of the largest and most well-studied communities of host-associated symbionts (2). The bacterial communities that colonize the gut and skin epithelia interact directly with potential pathogens in the environment, and can influence infection by stimulation of the host immune system, competition for space and nutrients, and production of inhibitory substances such as organic acids and bacteriocins (3–6). Interactions between the gut microbiota and pathogens of bees is an emerging area of research with both fundamental and applied importance (6, 7). Elucidation of the antipathogenic potential of the bee microbiota may ultimately help to preserve the pollination services provided by both wild and managed bees, which improve yields of over two-thirds of common agricultural crops (8) and contribute to the >$150B per year in economic value supplied by animal pollination (9).

The gut microbiota of corbiculate (“pollen basket”) bees, including honey and bumble bees, comprises a common core of five bacterial clades: *Snodgrassella* (Betaproteobacteria), *Gilliamella* (Gammaproteobacteria), *Bifidobacterium*, and *Lactobacillus* clades “Firmicutes-4” and “Firmicutes-5” (10). In addition to these core symbionts, bees may be infected by a variety of bacterial, fungal, protozoal, and viral pathogens (11), many of which are shared between wild and managed bees (12), can elevate mortality (13), and have been implicated in declines of bee populations on multiple continents (14–16). Both core and non-core microbiota have been found to stimulate immunity and enhance bee resistance to pathogens (6, 17–19). For example, depletion or perturbation of the gut microbiota increased the severity of bacterial, fungal, and protozoal infections in honey bees (6, 18, 20), whereas supplementation with core and hive-associated bacteria improved survival of infected larvae and adults (18, 21, 22). Several studies have shown direct inhibitory effects of gut and hive-associated symbionts against common bee pathogens (23–25), which suggests a parsimonious explanation for the effects of symbionts on infection.

The gut microbiota of bumble bees (*Bombus* spp.) has been repeatedly associated with resistance to infection with the trypanosomatid gut parasite *Crithidia bombi* (26–29). This parasite has a variety of negative effects on bees. Symptoms include reduced rates of foraging, pollen collection, and learning for worker bees (30–32); and reduced winter survival and spring nest-founding success for queen bees (33). Infection with *C. bombi* can also exacerbate susceptibility to co-occurring stressors such as starvation, pesticides (34), and nectar alkaloids (35). Trypanosomatid infections appear to be common in corbiculate bees, afflicting over half of individuals in some honey and bumble bee populations (36, 37), and have been correlated with honey bee colony collapses (37, 38) and native bumble bee declines (15). Both the presence and composition of the bumble bee microbiota may improve resistance to *C. bombi.* For example, germ-free rearing conditions and treatment with antibiotics both resulted in higher infection intensity in *B. terrestris* (26, 39). In contrast, absolute and relative abundance of select core gut bacteria were correlated with resistance to infection in both field surveys and fecal transplant experiments (26, 27, 29, 39). Just 3 taxa—*Snodgrassella, GIlliamella,* and *Lactobacillus* Firm-5*—*generally account for over 80% of the total gut bacteria in bumble bees (26, 29, 40). In both *B. terrestris* and *B. impatiens*, inoculation with microbiota rich in *Lactobacillus* Firm-5 resulted in resistance to *C. bombi* infection (26, 29). However, no study has examined the mechanisms by which these microbes influence parasite growth.

*Lactobacillus* Firm-5 is a group of lactic acid-producing bacteria (41). One way that they might increase host resistance to parasites is via modification of gut chemistry. Lactic acid fermentation results in production of organic acids that lower pH and inhibit growth of organisms that cause spoilage and infection (42–45). Indeed, lactic acid bacteria have a long history of use in food preservation in both human and insect societies (46, 47). In the host intestine, lactic acid bacteria can inhibit enteric pathogens, such as *Salmonella* and *E. coli* (48). This inhibition may reflect stimulation of the host immune system (49, 50), including that of insects (17, 23). However, *Lactobacillus-*mediated inhibition of pathogens is most simply explained by the direct antimicrobial activity of *Lactobacillus* metabolites. These metabolites, which include lactic acid and bacteriocins (45), may reduce the suitability of the gut environment for pathogens.

In the bee gut, *Lactobacillus* Firm-5 has been shown to have a disproportionately large effect on gut metabolomics. In honey bees, mono-inoculation with Firm-5 accounted for over 80% of the changes seen in bees inoculated with a full complement of gut microbes (51). Firm-5 isolates also showed *in vitro* inhibitory activity against the pathogens *Paenibacillus larvae* and *Melisococcus plutonius* (24). The high relative abundance of the Firm-5 clade in bumble bees (often >30% total bacteria (26, 40)), combined with its consistent association with resistance to trypanosomatid infection, suggests that *Lactobacillus* Firm-5 plays a major role in bumble bee resistance to trypanosomatid parasites.

We hypothesized that *Lactobacillus* Firm-5 enhances resistance to trypanosomatid parasites primarily by modifying the pH of the enteric environment. To test this hypothesis, we measured the inhibitory effects of spent medium from *Lactobacillus bombicola*, a member of the Firmicutes-5 clade that is ubiquitous in the gut microbiota of corbiculate bees, including *Bombus* spp. (10, 24, 40), on *in vitro* growth of several strains of *C. bombi.* We predicted that spent medium from *L. bombicola* would inhibit *C. bombi* growth, that the acidity of spent medium would be both necessary and sufficient to account for parasite inhibition, and that *C. bombi* strains would vary in sensitivity to spent medium.

## RESULTS

### Overview of experiments

Three experiments were conducted to evaluate effects of spent medium from *L. bombicola* cultures on growth of *C. bombi*. Spent medium was generated by growth of *L. bombicola* in MRS broth for 3 d, followed by sterile filtration to remove live cells. For *C. bombi* growth assays, the MRS-based spent medium (or MRS broth control) was diluted 1:1 in fresh, *Crithidia-*specific “FPFB” medium (52). The Neutralization Experiment tested whether spent medium would inhibit growth, and whether acidity of the spent medium was necessary or sufficient for inhibition. The Acidification Experiment tested for variation in pH-dependent growth inhibition due to various sources of acidity. The Strain Variation Experiment tested for variation in sensitivity to spent medium among different parasite strains.

### Neutralization Experiment: acidity-dependent inhibition of *C. bombi* by spent medium

We found that *L. bombicola* spent medium completely inhibited growth of *C. bombi* cell cultures (Figure 1). However, neutralized spent medium had no inhibitory effect, indicating that acidity of the spent medium was necessary for inhibition (Figure 1). Moreover, acidification of fresh (*Lactobacillus*-specific) MRS medium to pH 4.8 with L-lactic acid led to a level of growth inhibition that was comparable to that caused by pH 4.8 spent medium (Figure 1). This demonstrated that spent medium from *L. bombicola* inhibited *C. bombi* growth, and that changes in pH were necessary and qualitatively sufficient to account for this inhibition.

**Figure 1.**
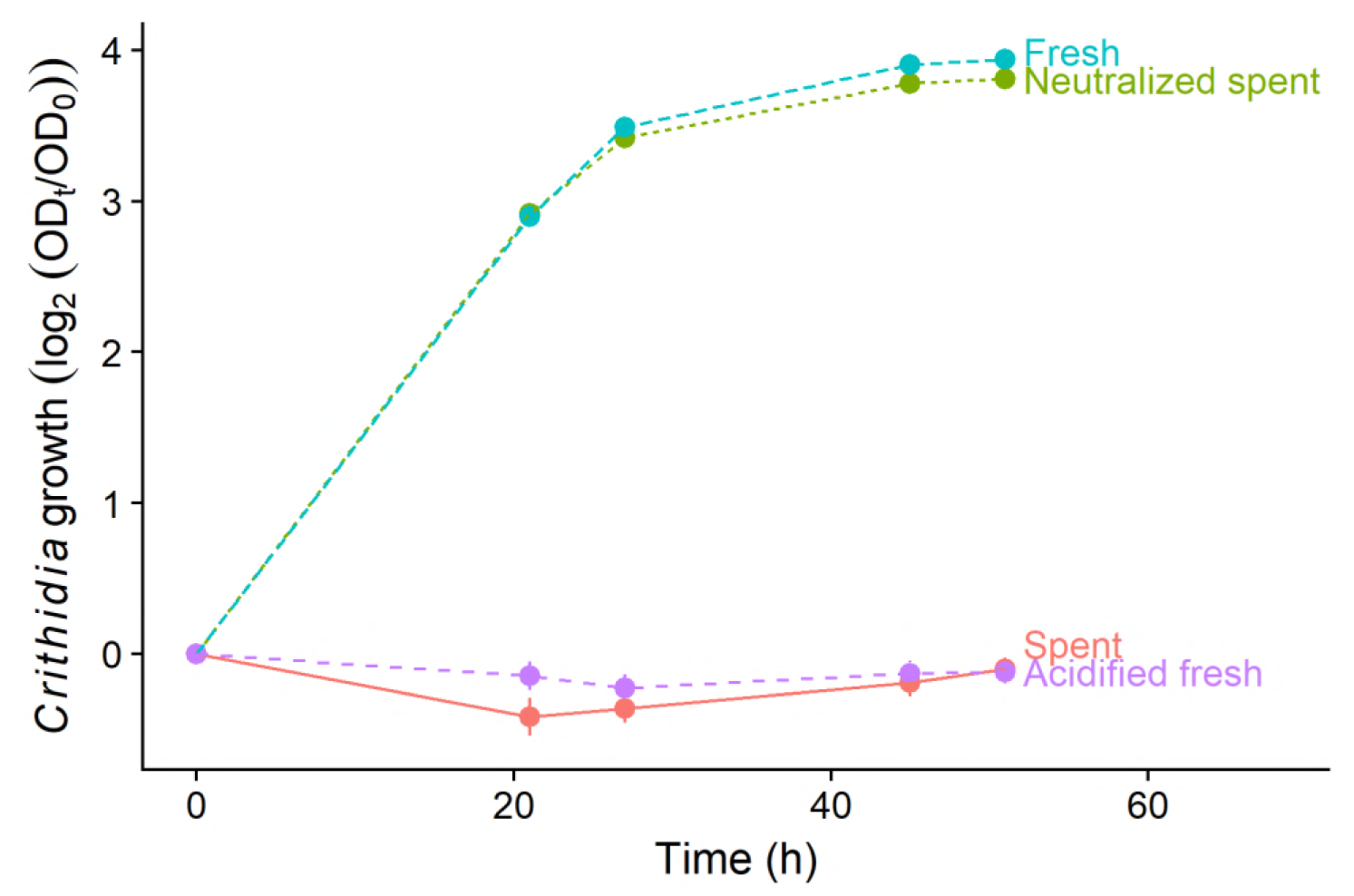
Spent medium from *Lactobacillus bombicola* inhibited growth of *Crithidia bombi;* acidity of the spent medium was necessary and sufficient for inhibition. Spent medium (“Spent”, red solid line) completely inhibited parasite growth. However, spent medium neutralized to pH 6.2 with NaOH (“Neutralized spent”, green dotted line) resulted in no inhibition relative to the fresh medium control (“Fresh”, light green dashed line), demonstrating that acidity was necessary for inhibition. Fresh medium acidified to pH 4.8 with lactic acid (“Acidified fresh”, blue dashed line) showed that acidity was sufficient for complete growth inhibition. Final pH of both spent medium and acidified fresh medium was 5.0 after combination with equal volume of fresh *Crithidia* medium (pH 5.9). Y-axis represents approximate number of parasite cell divisions, as measured by optical density (630 nm). Points and error bars show means and 95% confidence intervals (n = 36 wells per treatment).

### Acidification Experiment: pH-dependent inhibition of *C. bombi* with different sources of acidity

Having established pH-dependent growth inhibition, we conducted a follow-up experiment to quantify *C. bombi* growth rates across a range of pH values, and to compare the relative inhibitory effects of *L. bombicola* spent medium with that of three other sources of acidity: HCl, L-lactic acid (the form produced by animal cells (53)), and D-lactic acid (the form produced by *L. bombicola* (41)). We used log-logistic models to estimate and compare the EC50 pH (i.e., the pH that inhibited growth by 50% relative to the unacidified control treatment) for each source of acidity. All four sources of acidity resulted in qualitatively similar inhibition of *C. bombi* (Figure 2A), with considerable inhibition achieved within the pH range previously measured in honey bee guts (vertical lines in Figure 2A, from (54)). However, the EC50 pH varied somewhat across sources of acidity, with a hierarchy of pH-dependent inhibitory potency in the order HCl < L-lactic acid < D-lactic acid < spent medium (Figure 2B, see Supplementary Information Table S1 for table of EC50 values and Supplementary Data S1 for model parameters, confidence intervals on EC50 ratios, and raw data). An additional experiment with a second *C. bombi* strain (‘IL13.2’) confirmed the greater potency of spent medium relative to HCl-acidified medium across multiple parasite genotypes (Supplementary Figure 2).

**Figure 2.**
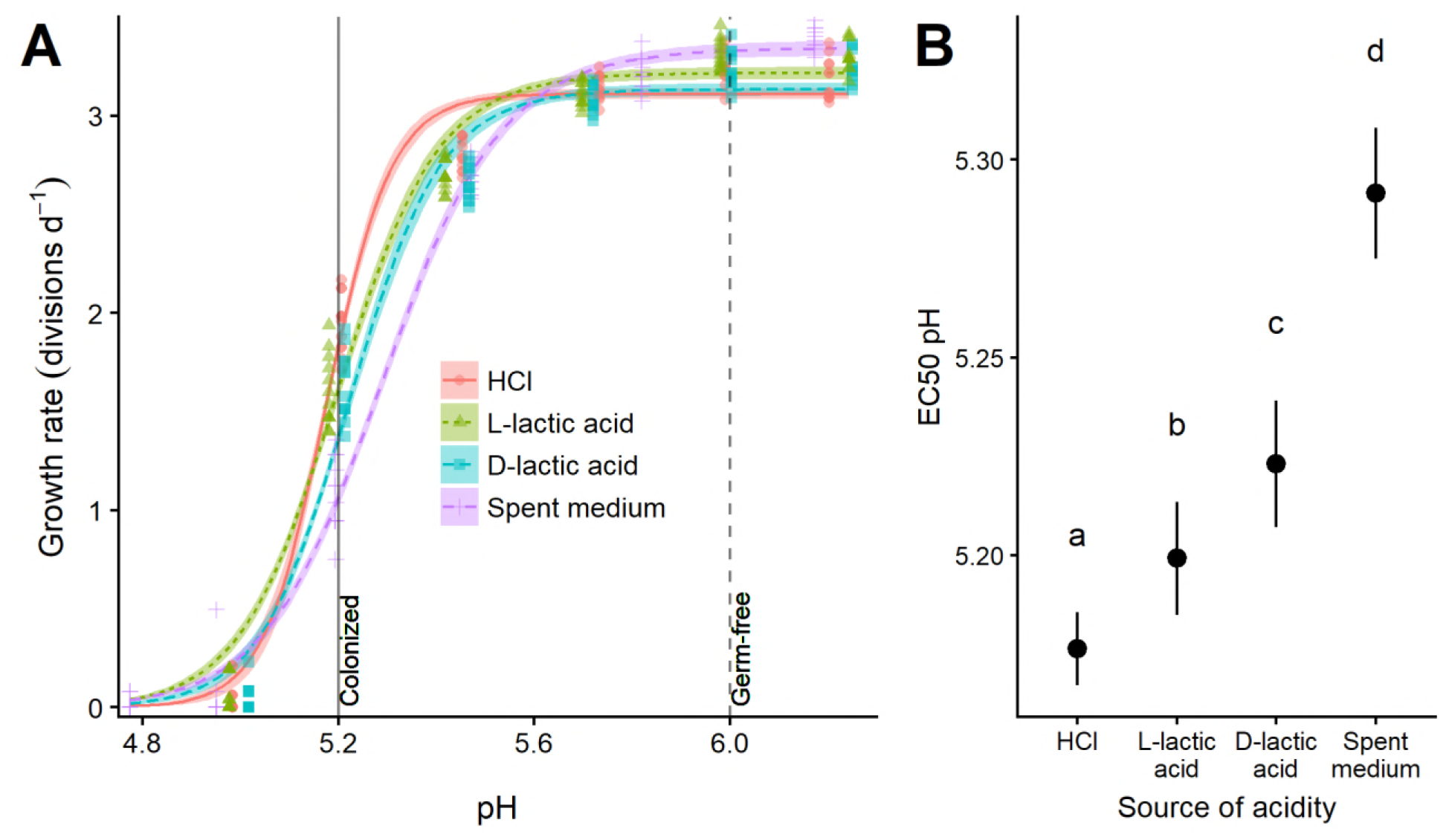
Different sources of acidity varied in pH-dependent inhibitory potency against *C. bombi.* (A) Dose-response curves relating pH to growth rate. X-axis shows final pH of treatment medium after combination of *L. bombicola* spent medium or acidified MRS medium with an equal volume of *Crithidia-* specific FPFB medium. Y-axis represents growth rate over first 21 h of incubation, measured as number of doublings per day by optical density (OD 630 nm). Lines and shaded bands represent model predictions and standard errors for each source of acidity. Points show raw data for each replicate well (n = 12). Vertical lines correspond to pH values measured in ileum and rectum of microbe-colonized and germ-free honey bees (54), and are shown as an estimate of gut pH range in bumble bees. **(B) 50% inhibitory concentrations** (EC50, i.e., the concentration that inhibits growth by 50%) for different sources of acidity. Point estimates and 95% confidence intervals are derived from model fits shown in panel **(A).** Higher EC50 pH estimates correspond to higher inhibitory potency for a given level of acidity. Different lower-case letters indicate statistically significant (P < 0.05) differences in pairwise comparisons of EC50 pH by source of acidity.

### Strain Variation Experiment: sensitivity to spent medium differs across *C. bombi* strains and rate of growth in culture (Figure 3)

We tested the dose-dependent effects of spent medium across three *C. bombi* strains, including two lines of strain VT1—the line that had been used for the above experiments and kept in continuous culture over the preceding 3 months (“VT1*), and a second line that had been thawed 20 d prior to the experiment (“VT1”). Values for EC50 pH showed statistically and biologically meaningful variation across cell lines (Figure 3). The line most thoroughly acclimated to the culturing conditions, VT1*, exhibited both the fastest growth (Figure 3A) and the lowest sensitivity to spent medium, as indicated by its low EC50 pH value (Figure 3B); strain IL13.2 exhibited the second-fastest growth and the second-lowest sensitivity to spent medium. Strains VT1 and S08.1 had similar growth rates in the absence of spent medium (Figure 3A), but had significantly different EC50 pH values (Figure 3B; see Supplementary Information Table S1 for table of EC50 values and Supplementary Data S1 for model parameters, confidence intervals on EC50 ratios, and raw data). All *C. bombi* strains suffered growth inhibition within the pH range (5.2-6.0) documented in the honey bee hindgut (54) (Figure 3A).

**Figure 3.**
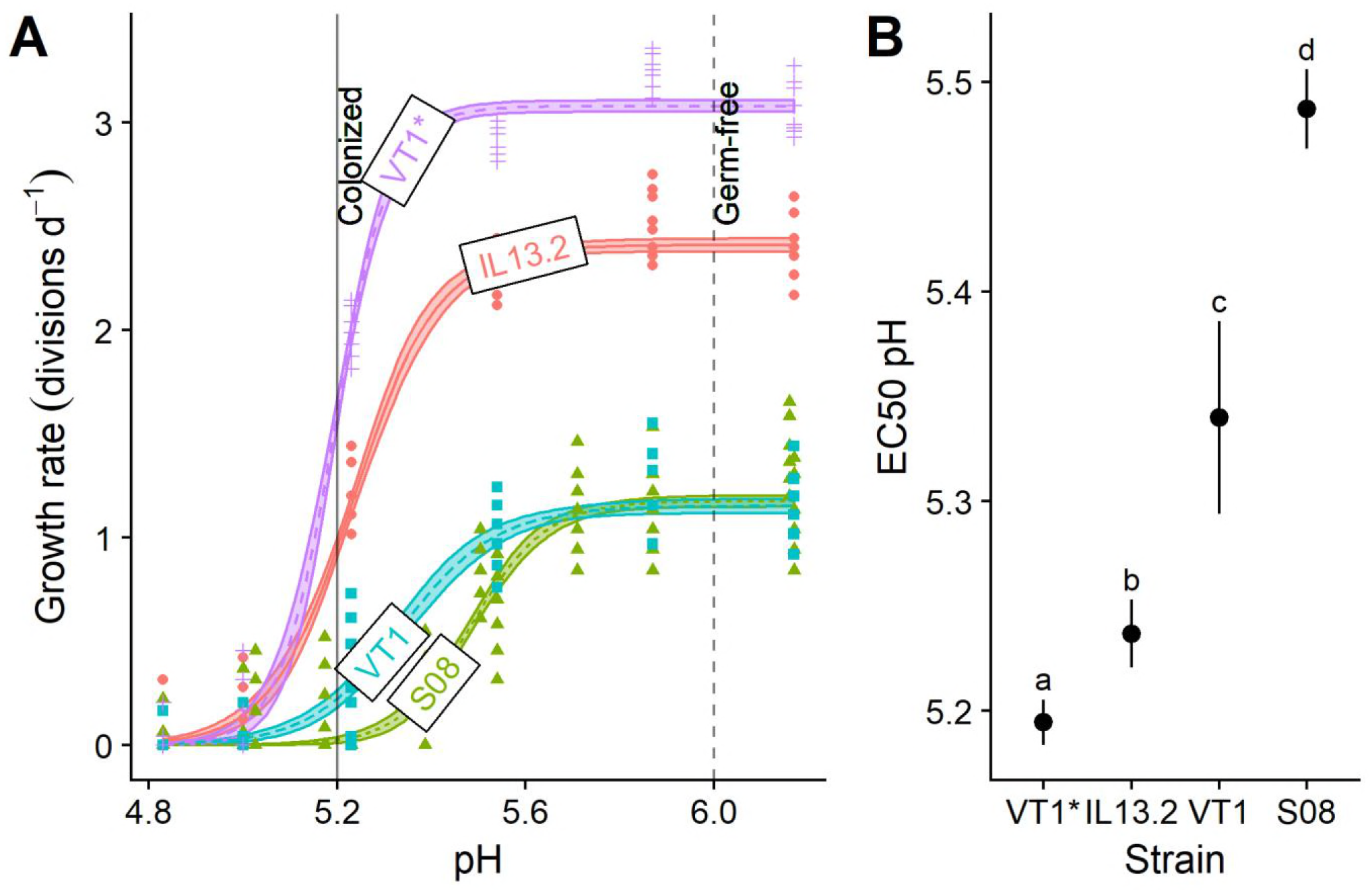
*Crithidia bombi* pH sensitivity varied according to strain identity and rate of growth in culture. (A) Dose-response curves relating pH to growth rate for different *C. bombi strains*. X-axis shows final pH of treatment medium after combination of *L. bombicola* spent MRS medium with an equal volume of *Crithidia-*specific FPFB medium. Y-axis represents growth rate over first 20 h of incubation, measured as number of doublings per day by optical density (OD 630 nm). Lines and shaded bands represent model predictions and standard errors for each strain. Points show raw data for each replicate well (n = 12). Vertical lines correspond to pH values measured in ileum and rectum of microbe-colonized and germ-free honey bees (54), and are shown as an estimate of gut pH range in bumble bees. Strains “VT1*” and “VT1” are the same strain, but “VT1*” had been grown in continuous culture for 3 months prior to the experiment, whereas “VT1” and all other strains had been thawed frozen cryopreserved stock 3 weeks prior. **(B) 50% inhibitory concentrations** (EC50) for each strain. Point estimates and 95% confidence intervals are derived from model fits shown in panel **(A).** Higher EC50 pH estimates correspond to higher sensitivity to acidity. Different lower-case letters indicate statistically significant (P < 0.05) differences in pairwise comparisons of EC50 pH by source of acidity. For all strains, inhibition occurred within the pH range measured in honey bee guts (data from (54)).

## DISCUSSION

Our results show that production of acids by the core bumble bee hindgut symbiont, *L. bombicola,* is both necessary and sufficient to inhibit growth of the widespread trypanosomatid parasite *C. bombi*. The growth-inhibitory effects of *L. bombicola*-acidified spent medium were qualitatively similar across *C. bombi* isolates and occurred within a pH range that is physiologically realistic for the bee hindgut where *C. bombi* establishes. Because Lactobacilli and other acid-producing bacteria are dominant members of the bumble bee gut microbiota, our results suggest that the inhibitory effects of bumble bee gut microbiota on *C. bombi* infection intensity can be largely attributed to microbial acidification of the gut. Our findings provide a mechanistic basis to understand how microbiota may affect trypanosomatid infection in corbiculate bees that share a core microbiome (10) and can be infected with identical and related parasites, including trypanosomatids (55–57).

### Inhibitory activity of *L. bombicola* spent medium is driven by production of acids

The effects of *L. bombicola* spent medium on *C. bombi* growth could be explained by the acidity of the spent medium. Environmental pH is a recognized driver of interactions in microbial communities, where species vary in how they alter the pH of their surroundings and in the pH range at which they can grow (58, 59). In honey bees, microbial colonization with core symbionts resulted in acidification of the hindgut lumen (54). In our cell culture experiments, a reduction in pH that corresponds to the difference between guts of germ-free (pH ̃6.0) and normal, hive-reared honey bees (pH ̃5.2 (54)) profoundly inhibited growth of *C. bombi*. Our findings indicate that symbiont-mediated gut acidification could act as an important filter that prevents trypanosomatid establishment and defends bees against infection.

### Spent medium was more strongly inhibitory than expected based on pH alone

Whereas all four tested acids inhibited *C. bombi* in a dose-dependent fashion, the EC50 pH varied across sources of acidity (Figure 2). Both enantiomers of lactic acid were more inhibitory than HCl for a given pH. This finding is consistent with prior work on bacteria, which showed growth inhibition at relatively high pH when lactic acid, rather than HCl, is used as the acidulant (42, 43). For example, the inhibitory pH of *E. coli* was 0.1 pH units higher with lactic acid, rather than HCl, as the acidulant (43). This increased activity reflects the fact that lactic acid and other weak acids are undissociated at low pH, which allows them to penetrate membranes of target cells with relative ease. Once inside the relatively alkaline cytoplasm of the cell, the acid dissociates, and disrupts the proton motive force necessary for energy production and homeostasis (44).

We also found very slightly but significantly higher potency of D-lactic acid relative to L-lactic acid (Figure 2). L-lactic acid is the enantiomer more often produced by trypanosomatids, including by *Leishmania* species that are closely related to *C. bombi* (60), and L-lactic acid dehydrogenase has been found in the genome of honey bee-infective trypanosomatids (61). It is possible that *C. bombi* can use oxidative phosphorylation (60) to metabolize this enantiomer, and thereby reduce its toxicity, or that they have greater resistance to the enantiomer that they produce through their own metabolism. For example, *L. bulgaricus* produced D-lactic acid, and were less inhibited by this self-generated enantiomer than by L-lactic acid (62).

Prior measurements of the pH range and sources of acidity in honey bee gut both indicate that pH-mediated inhibition of trypanosomatids is achievable for corbiculate bees. The pH of a symbiont-colonized honey bee hindgut (̃5.2 (54)) was similar to the EC50 pH for HCl (5.18) and lower than the EC50 pH due to spent medium (5.29). *In vitro, L. bombicola* produced exclusively D-lactic acid (53), which was the most inhibitory (i.e., highest EC50 pH) of the pure acids tested here (Fig 3). Moreover, the specific acids that are found in the gut environment are generally more inhibitory for a given pH than those tested here. In the honey bee hindgut, the most abundant acids were acetic acid and succinic acid (54). Both acetic acid (pKa = 4.95) and succinic acid (pKa = 4.2 for first deprotonation and 5.6 for second deprotonation) have relatively high pKa’s relative to lactic acid (pKa = 3.86) and HCl (pKa = −7). These high pKa values mean that at low pH, the acids are found in their more potent, undissociated form. As a result, these high-pKa acids generally have antimicrobial effects that are even stronger than those of lactic acid for a given pH (42). Indeed, lactic and acetic acids can have synergistic effects against growth of *E. coli* (42), with lactic acid producing a low pH that increases the fraction of undissociated acetic acid (42).

Spent medium was slightly more inhibitory than all pure acids for a given pH (Figure 3). Because neutralization completely removed the inhibitory activity of spent medium, it appears that some aspect of the spent medium—whether the existence of some metabolite or the relative lack of nutrients—is only inhibitory at low pH. In other words, something about the spent medium is potentiated by an acidic environment. For example, low pH could facilitate solubility or penetration of non-lactic acid components, such as bacteriocins; this type of synergy was seen in other studies of *Lactobacillus* spp. (63, 64). Within the bee gut, synergistic effects could also occur between organic acids and toxins produced by other members of the microbiota (41, 65).

### Strains varied in sensitivity to spent medium

We found that sensitivity to spent medium varied by *C. bombi* strain and rate of growth in culture. Strain VT1 that had been in continuous culture for 3 months and had the fastest growth rate was the least sensitive to spent medium, followed by the next-fastest strain IL13.2, the recently thawed line of Strain VT1, and strain S08.1. The comparison between the recently thawed VT1 and S08.1 strains—which had similar maximal growth rates, but different levels of sensitivity to spent medium—indicates that differences in pH sensitivity have a genotypic as well as environmental basis and are not purely driven by the overall growth rate in culture. Strains of *C. bombi* have been shown to be both genetically and phenotypically diverse, and to vary in growth rate (66), infectivity (67), and responses to host diet composition (68), phytochemicals (69), and microbiota (26). Our study documents variation in pH sensitivity within an ecologically relevant range of pH that is likely representative of the environment in the bumble bee gut.

Our results showed some growth inhibition of all strains—and complete inhibition of some strains—within the pH range measured in the gut of honey bees (54). The pH found in a germ-free gut (5.8-6.0) was favorable for growth, consistent with the high *C. bombi* infection intensities found in germ-free and antibiotic-treated bees (39), whereas pH of a symbiont-colonized gut (<5.2) would be expected to inhibit growth of all strains. Thus, pH sensitivity may constrain ability of strains to colonize certain host genotypes (67) or enterotypes (26, 70) characterized by low gut pH. For example, inoculation with a microbiota high in *Lactobacillus* Firm-5 resulted in lower overall infection intensity and favored infection with a single parasite strain that was less successful in bees inoculated with microbiota low in Firm-5 (26). We hypothesize that low-pH conditions favor strains that are more tolerant, whereas high-pH conditions favor strains that are strong competitors. Further experiments are needed to investigate the extent to which parasite populations are selected for pH tolerance, and possible trade-offs between growth rate, infectivity, or tolerance to environmental stressors and insect immune factors. Environment-dependent selection for these other traits could maintain variation in pH tolerance within parasite populations.

### Gut microbiota-driven changes in pH may explain patterns of trypanosomatid infection in bees

The pH-dependent inhibition demonstrated here is consistent with past surveys and experiments that showed negative correlations between abundance of acid-producing gut symbionts and *C. bombi* infection intensity, and with associations between microbiota composition and relative infectivity of different parasite strains. The *Bombus* gut microbiota is dominated by three taxa—*Lactobacillus* Firm-5, *Gilliamella, and Snodgrassella,* that made up over 80% of total gut bacteria (26, 40). In fecal transplant experiments *of B. terrestris* and *B. impatiens,* high relative abundance of *Lactobacillus* Firm-5 and *Gilliamella* were negatively correlated with *C. bombi* infection intensity (26, 28, 29, 39). Both *Lactobacillus* and *Gilliamella* ferment sugars to produce acids. *Lactobacillus* Firm-5 made a particularly strong contribution to the gut chemical profile in honey bees (51). *Gilliamella apicola* produced acids from diversity of sugar sources, including sugars that are toxic to honey bees (71), and produces metabolites that feed the co-localized core bacterium *S. alvi* (51).

In contrast, *Snodgrassella* abundance was not correlated with resistance to *C. bombi* in either *B. terrestris* or *B. impatiens* (26, 39). On the contrary, pre-inoculation of honey bees with *S. alvi* resulted in gut microbial overgrowth and increased trypanosomatid infection intensity (70). Whereas *Lactobacillus* and *Gilliamella* spp. produce acids from sugars, *Snodgrassella* consumes organic acids (51, 72). This metabolic activity could elevate gut pH to levels that are more hospitable to trypanosomatids. On the other hand, *Snodgrassella* abundance was negatively correlated with infection prevalence in a field survey (27). This correlation might reflect the association of *Snodgrassella* with acid-producing *Gilliamella* (51), rather than inhibitory effects of *Snodgrassella per se*. *Snodgrassella* and *Gilliamella* form a biofilm that lines the ileum (71) and could competitively inhibit trypanosomatid attachment to the gut wall (55, 73). Further study would be needed to determine the relative contributions of gut acidification versus biofilm formation to microbiota-induced inhibition of parasites in bee guts. It would be intriguing to investigate how *Snodgrassella* abundance alters gut pH, and whether the effects of this symbiont on gut pH are outweighed by its direct competition with trypanosomatids for space along the gut wall, or by its contribution to anoxic gut environments (54) that can be toxic to insect gut trypanosomatids (60). Because *Apis* and *Bombus* share a core gut microbiota (74, 75), we hypothesize that trypanosomatid parasites of honey bees (55) and bumble bees interact with similar communities of gut symbionts, and that the same pH-altering symbiont taxa could govern parasite establishment in both host genera.

### Do gut microbiota shape resistance to opportunistic infection via alteration of gut pH?

In the context of prior experiments, our results suggest a pH-mediated role of the bee microbial community, both in the hive and the gut itself, in defense against opportunistic infection. Multiple studies have correlated lack of core gut bacteria, or abundance of non-core bacteria, with bee infection. In North America and Europe, high diversity of non-core bacterial species correlated with higher prevalence of the pathogens *Crithidia* and *Nosema* (27, 28). Similarly, in a survey of *Bombus* spp. in China, bees could be grouped into two microbial enterotypes (70). One of these enterotypes was less variable and dominated by core symbionts *Snodgrassella* and *Gilliamella.* In contrast, the other enterotype was highly variable and comprised non-core *Hafnia*, Enterobacteriaceae, and *Serratia*, The latter two are thought to be opportunistic pathogens (6, 16). Finally, higher susceptibility to pathogens was found in germ-free honey and bumble bees (6, 18, 29, 39). The acidification of the bee gut lumen by symbionts could explain how a strong core microbiota resists invasion by pathogens. This hypothesis is consistent with the dominant role of pH in determination of bacterial communities in soil (58, 59), and correlations between low gastric acidity and opportunistic infection in human subjects (76). However, other explanations for the effects of core gut microbiota on parasites—such as physical competition for space and resources, enhancement of the immune response, and improvement of host nutritional status (6)—also deserve consideration. It is possible that different microbial assemblages are optimal for different functions. For example, a microbiome that produces abundant acids might be optimal for direct defense against parasites, whereas other community compositions might be provide more benefits to immune system regulation, or to nutritional sufficiency that improves resistance or tolerance to infection (33, 77). Further research is needed to investigate variation in gut pH within and across species, and possible causative relationships between microbiome composition and bee health under different environmental circumstances.

### Conclusion

Our results build on prior associations between microbiome and infection intensity to provide mechanistic insights into how the bee gut microbiota are likely to influence trypanosomatid infection in bumble bees and possibly other corbiculate bee species. We have shown that symbiont-mediated reductions in pH constrain parasite growth in culture; that inhibitory pH in vitro varies according to parasite strain and level of acclimation to culture conditions; and that the range of variation in inhibitory pH was within the range of pH measured in the honey bee gut (54). These findings suggest that the pH of the gut environment may be a critical determinant of trypanosomatid growth *in vivo*, a hypothesis that could be tested in both manipulative and observational studies that relate gut pH to trypanosomatid establishment. The role of gut pH in resistance to other enteric pathogens, and the selective forces acting on symbionts and parasites to create and tolerate different levels of acidity, warrants further study as part of continued investigations into the functional significance of the bee microbiome.

## MATERIALS AND METHODS

### Cell Cultures

*Lactobacillus bombicola* strain 70-3, isolated from *Bombus lapidarius* collected near Ghent Belgium (isolate “28288T” (53)), was obtained from the DSMZ and grown in 2 mL screw-cap tubes in MRS broth (Research Products International, Mt. Prospect, IL) with 0.05% cysteine at 27 °C. *Crithidia bombi* cell cultures were isolated from bumble bee intestines by flow cytometry-based single cell sorting as described previously (52). Cultures originated from wild infected bumble bees. Strains VT1 (Vermont, USA, 2013, courtesy Rebecca Irwin) and IL13.2 (Illinois, USA, 2013, courtesy Ben Sadd) originated from *B. impatiens* workers. Strains S08.1 (Switzerland, 2008, courtesy Ben Sadd) originated from *B. terrestris.* These same cell lines have been used to assess effects of phytochemicals on parasite growth (78). Briefly, cells from fecal samples were sorted into 96-well plates containing “FPFB” culture medium with 10% heat-inactivated fetal bovine serum and incubated at 27 °C. Cultures with successful growth and absence of visible contamination were transferred to vented, 25 cm^2^ tissue culture flasks, grown to high density, and cryopreserved at −80 °C until several weeks before the experiments began (52). Culture identity was confirmed as *C. bombi* based on glyceraldehyde 3-phosphate dehydrogenase and cytochrome b gene sequences.

The Neutralization and Acidification Experiments were performed with a line of strain “VT1” that had been in continuous culture for 2 months at the start of the experiments presented here, with transfers to fresh medium every 3-4 d. This line is referred to as “VT1*” in the Multi-Strain Experiment, to differentiate it from the more recently thawed line of the same strain. All other strains in the Multi-Strain Experiment were thawed 20 d prior to the assay and transferred to fresh medium every 3-7 d, depending on growth.

### Generation of spent medium

To generate spent medium, *L. bombicola* aliquots were transferred to 8 mL fresh medium and grown in screw-cap 14 mL conical tubes at 27 °C for 3 d. The resulting spent medium (net OD_630 nm_ = 0.500-0.700) was sterile-filtered through a 0.22 µm membrane and stored at −20 °C until use in experiments (not more than 2 weeks). Fresh MRS medium, incubated under identical conditions, was used as a control.

### Neutralization Experiment

To evaluate the inhibitory effects of spent medium, neutralized spent medium, and acidified fresh medium, growth *of C. bombi* was compared across four MRS-based treatments: Spent medium (“Spent”, initial pH 4.8), spent medium neutralized to pH 6.2 with 1 M NaOH (“Neutralized spent”), fresh MRS medium (“Fresh”, pH 6.2), and fresh MRS medium acidified to pH 4.8 with 1 M HCl (“Acidified fresh”). The *C. bombi* culture was diluted to 2x desired final concentration in fresh FPFB medium (pH 5.88). The resulting cell suspension (100 µL) was added to wells of a 96-well plate containing an equal volume of the MRS-based treatment medium, resulting in a final net OD_630 nm_ of 0.010. *Crithidia* cultures were found to grow adequately in mixed medium, but *L. bombicola* failed to grow in 100% FPFB medium, and *C. bombi* failed to grow in 100% MRS medium. Growth was measured twice daily by optical density over the ensuing 48 h. Net optical density at each time point was computed by subtracting OD of wells containing the corresponding MRS-based treatment medium and FPFB medium without *C. bombi*; this controlled for any differences in optical density that occurred independent of *C. bombi* growth. The experiment included 18 replicate wells per treatment.

### Acidification Experiment

To compare variation in growth inhibition across different sources of acidity, growth of *C. bombi* was compared in dilutions of spent medium (initial pH 4.65), and in fresh medium acidified with D-lactic acid (pH 4.82), L-lactic acid (pH 4.77), or HCl (pH 4.73). Each base medium was diluted with fresh MRS medium to 0, 20, 40, 60, 80, or 100% of initial concentration. The final pH of each treatment was measured with a pH meter (‘Orion Star’, Thermo, Waltham, MA) after combination with an equal volume of FPFB medium. Growth of *C. bombi* (initial OD 0.010) was evaluated with a 96-well plate assay as in the Neutralization Experiment above. The experiment included 12 replicate wells per concentration of each acidification treatment. To verify the relative potency of spent vs. acidified medium against different parasite strains, the experiment was repeated with two *C. bombi* strains (IL13.2 and VT1*) tested in parallel against two of the four acidification treatments (spent medium and HCl-acidified medium); results are shown in Supplementary Figure 2.

### Strain Variation Experiment

To compare susceptibility to spent medium across different *C. bombi* strains and degrees of acclimation to the culture environment, inhibitory concentration of spent medium was compared across four cell lines: the “VT1*” line of strain VT1 that had been used for the above experiments, and by this time had been in continuous culture for 3 months; and strains VT1, IL13.2, and S08.1 that had been thawed 20 d prior to the experiment. Six dilutions of spent medium (0-100%) were prepared in fresh MRS medium; final pH of each treatment was measured after combination with an equal volume of FPFB medium. Growth of each line of *C. bombi* (initial OD 0.010) was evaluated with a 96-well plate assay as in the Neutralization Experiment above. The experiment included 12 replicate wells per cell line and spent medium concentration.

### Statistical analysis

Statistical analysis was performed in R v3.4.3 for Windows (79). For the Acidification and Strain Variation Experiments, dose-response curves to relate growth rate to pH were estimated with the *drm* function in R package *drc* (80). Under the conditions of the experiment (initial OD = 0.010, incubation temperature 32 °C), *Crithidia bombi* growth was found to be exponential over the first 24 h of incubation (Supplementary Figure 1). Therefore, the rate of growth (log_2_(OD/OD_0_)/t) over the first incubation interval (0-21 h) was used as the response variable for the dose-response models. A log-logistic function, with the lower asymptote fixed at 0, was fitted to the growth measurements for each acidification treatment (Acidification Experiment) or cell line (Strain Variation Experiment), using final pH of the treatment medium at the start of the experiment (i.e., after combination of the MRS-based treatment with an equal volume FPFB medium) as the predictor variable. The 95% confidence intervals for EC50 pH (i.e., the pH that inhibited growth by 50% relative to the unacidified control treatment), and 95% CI’s for ratios of EC50 pH among acidification treatments and strains, were estimated using the delta method (*drc* function *EDcomp* (80)). Acidification treatments and strains were considered significantly different when the 95% confidence interval for the ratio of their EC50 pH values did not include 1.

## ACKNOWLEDGMENTS

The authors thank Ben Sadd for providing strains IL13.2 and S08.1, Rebecca Irwin for providing the bees from which Strain VT1 was established, and Guang Xu and Ben Sadd for sharing DNA sequences. This project was funded by a National Science Foundation Postdoctoral Research Fellowship to EPY (NSF-DBI-1708945); USDA NIFA Hatch funds (CA-R-ENT-5109-H), NIH (5R01GM122060-02), and NSF MSB-ECA (1638728) to QSM; and an NSF-CAREER grant (IOS 1651888) to TRR. The funders had no role in study design, data collection and interpretation, or the decision to submit the work for publication. The authors declare that they have no conflicts of interest.

## DATA AVAILABILITY

All data are supplied in the Supplementary Information, Data S1.

